## supplementary files for "Propagation of Hippocampal Ripples to the Neocortex by Way of a Subiculum-Retrosplenial Pathway"

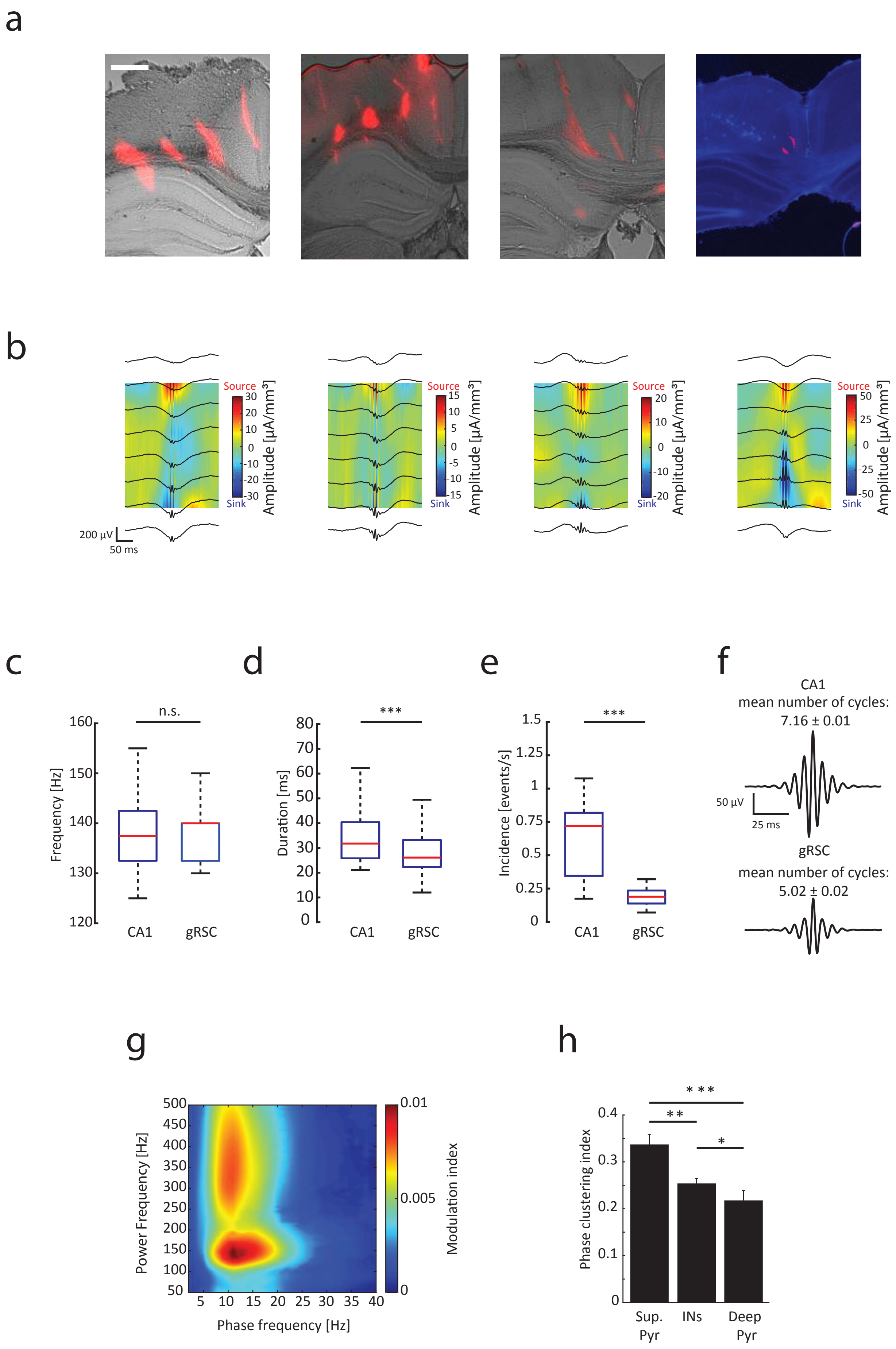
a
b
Source
Sink
Amplitude [μA/mm³]
200 μV
50 ms
c
d
e
f
n.s.
Frequency [Hz]
CA1
gRSC
***
Duration [ms]
Incidence [events/s]
CA1
mean number of cycles:
7.16 ± 0.01
50 μV
25 ms
gRSC
mean number of cycles:
5.02 ± 0.02
g
h
Power Frequency [Hz]
Phase frequency [Hz]
Modulation index
Phase clustering index
Sup. Pyr
INs
Deep Pyr

**Supplementary Figure 1. Characterization of gRSC ripple activity.** a) Histological verification of recording location. Silicon probes were coated with DiI (red) and the location of individual recording sites was estimated based on recording depth and reconstructed probe tracks. Shown are 4 out of 6 animals where superficial layers were successfully targeted. For the remaining examples see Fig. 1a and Fig. 2a. Scale bar, 500 µm. b) CSD maps centered around gRSC ripple peak for the remaining subjects not shown in Fig. 1d. c) Statistics of ripple frequency in CA1 (left) and gRSC (right). Red lines, median; blue lines, quartiles. n.s., not significant. d) Statistics of CA1 (left) and gRSC (right) ripple duration ($p<0.001$, rank sum test). e) Statistics of CA1 (left) and gRSC (right) ripple incidence ($p<0.001$, rank sum test). f) Average waveforms and number of cycles of CA1 (top) and gRSC (bottom) ripples (n = 19 sessions from 6 animals; 22,694 CA1 ripples, 8,501 gRSC ripples). g) Averaged phase-amplitude comodulogram for superficial gRSC LFP showing cross-frequency coupling between gRSC ripples and negative waves occurring at around 10 Hz (n = 10 sessions from 6 mice, $p<0.05$ shuffling analysis with 200 shuffles). h) Phase clustering index (mean ± s.e.m) of all significantly phase modulated superficial pyramids, deep pyramids and interneurons (superficial pyramids ITPC = 0.34 ± 0.02; deep pyramids ITPC = 0.14 ± 0.03; INs ITPC = 0.21 ± 0.01; ***$p < 0.001$, **$p<0.01$, *$p<0.05$).

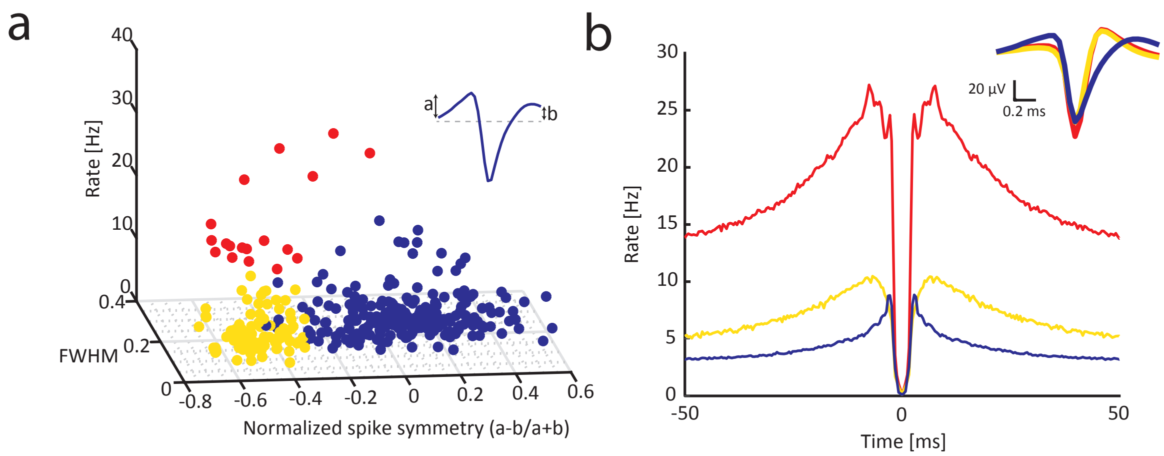


**Supplementary Figure 2. Cell type classification.** a) Units were separated into putative fast-spiking INs (red), low-spiking INs (yellow) and pyramids (blue) based on firing rate, waveform width (FWHM) and waveform asymmetry (inset). Low and fast spiking INs were re-grouped. b) Average autocorrelograms for the three groups (inset, average waveforms), based on n = 335 units recorded from all layers from n = 8 mice.

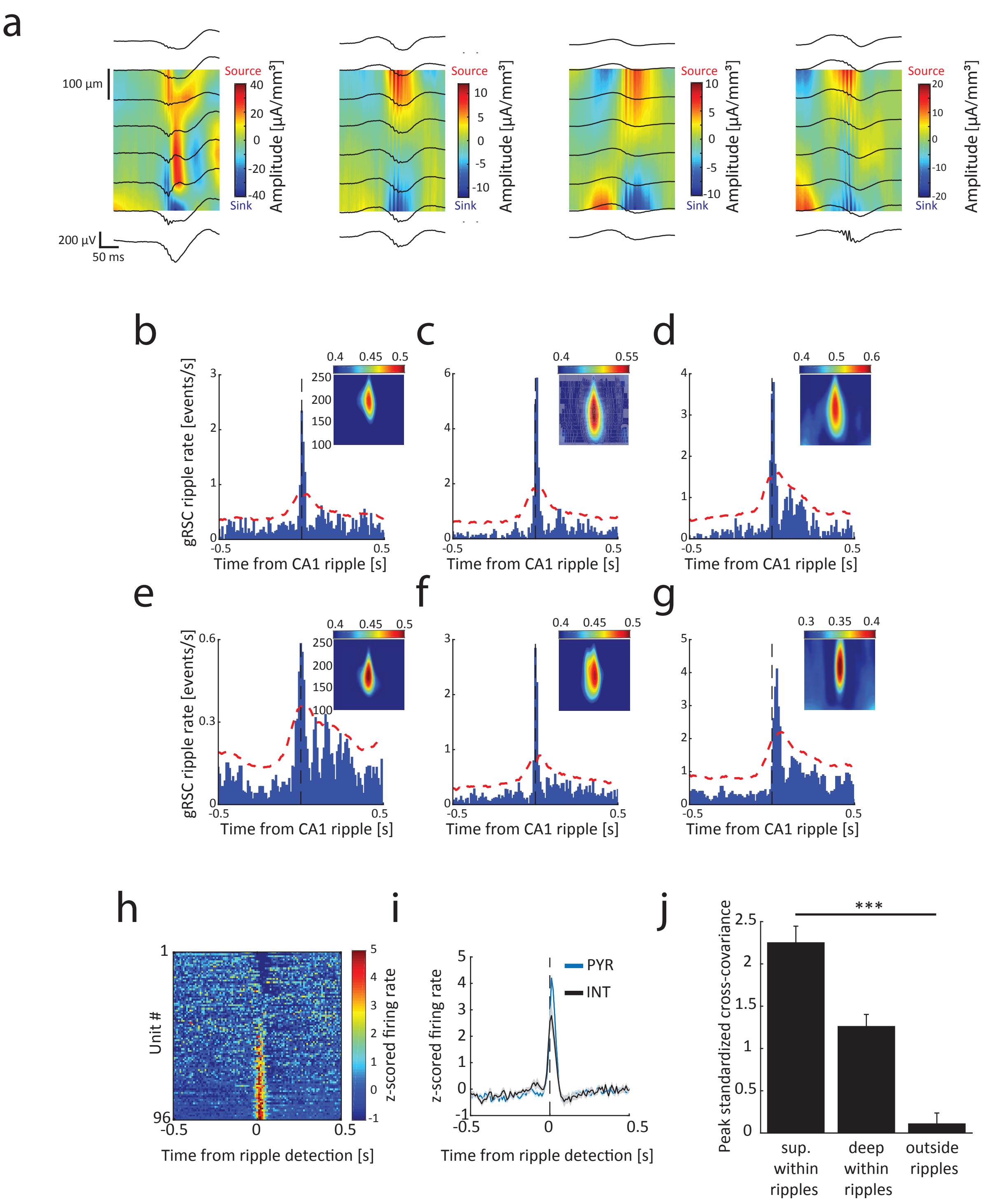


**Supplementary Figure 3. Coupling between gRSC ripples and CA1 SPW-Rs.** a) gRSC CSD maps centered around CA1 SPW-R peak for the remaining subjects not shown in Fig. 2h. b-g) CA1-gRSC ripple CCGs and power coherograms (inset, y-axis range 100 to 250 Hz, x-axis range ± 50 ms) for six individual animals. Black lines, CA1 SPW-R detection (b = 5 sessions 3,555 CA1 and 1,623 gRSC ripples; c = 2 sessions 2,170 CA1 and 513 gRSC ripples; d = 4 sessions, 2,119 CA1 and 1,157 gRSC ripples; e = 2 sessions, 2,106 CA1 and 710 gRSC ripples; f = 4 sessions, 7,998 CA1, 1,244 gRSC ripples; g = 2 sessions 4,746 CA1, 3,254 gRSC ripples). Red lines, upper 95% confidence interval. h) Z-score normalized PETH showing the responses of CA1 and subiculum pyramidal units to SPW-Rs sorted by response magnitude. i) Averaged z-scored responses of hippocampal units separated into putative pyramidal cells (blue) and INs (grey). j) Peak standardized cross-covariance (mean ± s.e.m) of superficial and deep gRSC units within ripple epochs (± 250 ms from ripple detection) and of units from all layers outside of ripple epochs. All groups are significantly different from each other ($p<0.001$ Bonferroni-corrected rank sum tests).

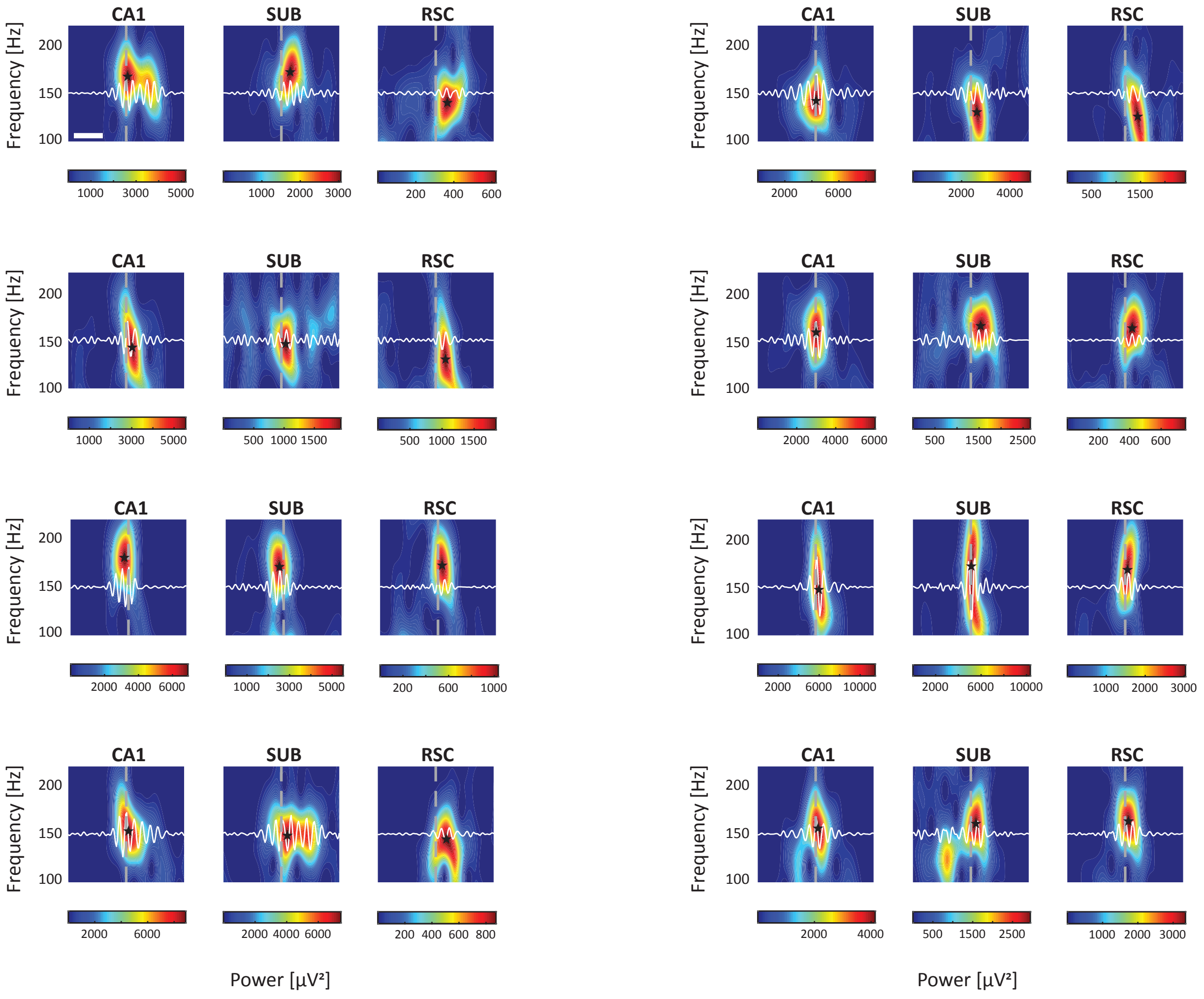


**Supplementary Figure 4. Propagation of ripple activity along the hippocampal output axes.** Individual examples of simultaneously detected CA1, subicular (SUB) and gRSC ripples. Ripple band filtered traces are overlaid in white. Dashed grey line denotes maximal amplitude ripple wave detected in CA1. Black star denotes ripple peak power. Scale bar, 25 ms.

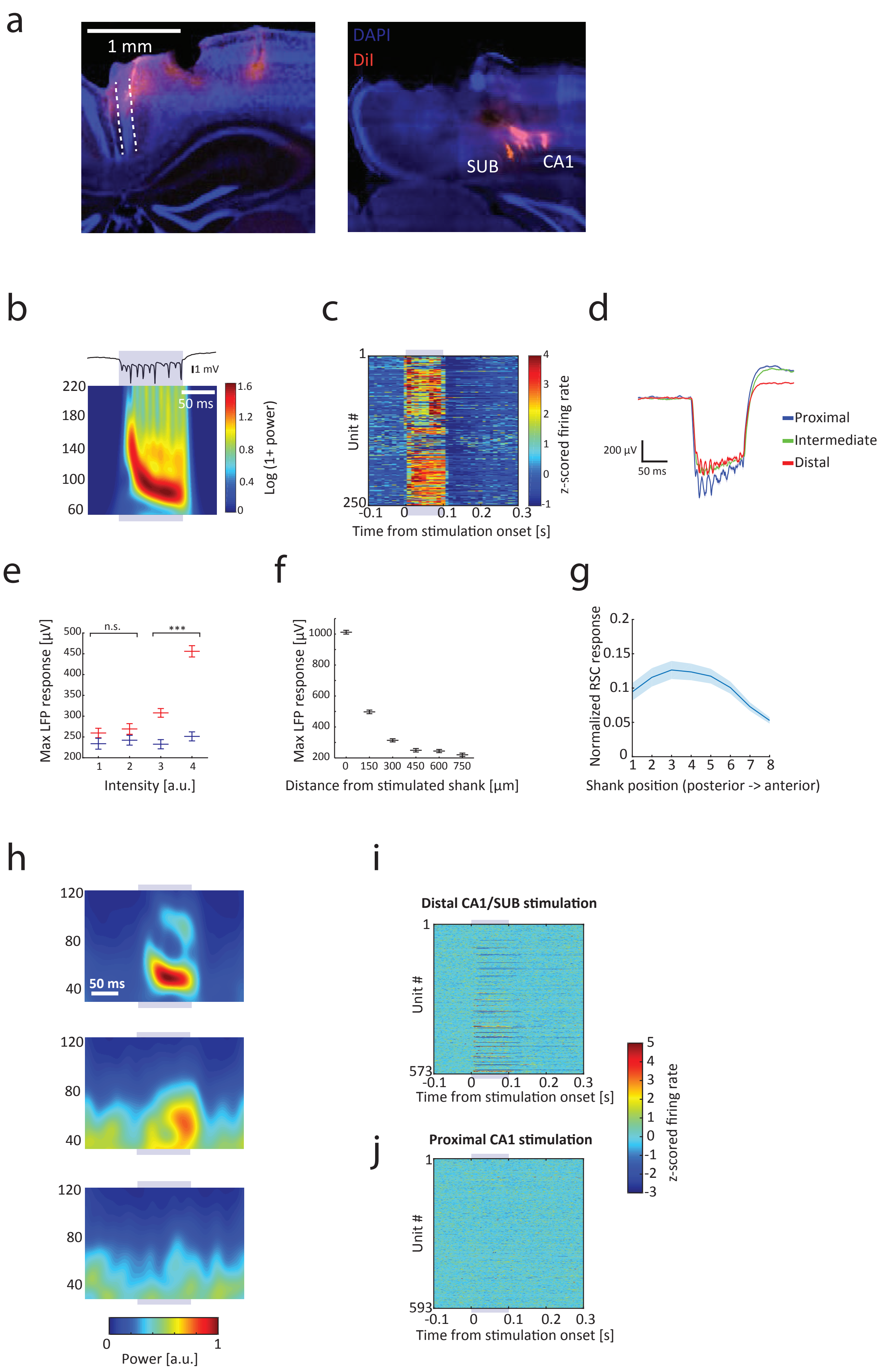
a
1 mm
DAPI
DiI
SUB
CA1
b
1 mV
50 ms
220
180
140
100
60
1.6
1.2
0.8
0.4
0
Log (1+ power)
c
1
250
Unit #
4
3
2
1
0
-1
z-scored firing rate
-0.1
0
0.1
0.2
0.3
Time from stimulation onset [s]
d
200 µV
50 ms
Proximal
Intermediate
Distal
e
Max LFP response [µV]
500
450
400
350
300
250
200
n.s.
***
1
2
3
4
Intensity [a.u.]
f
Max LFP response [µV]
1000
800
600
400
200
0
150
300
450
600
750
Distance from stimulated shank [µm]
g
Normalized RSC response
0.2
0.15
0.1
0.05
0
1
2
3
4
5
6
7
8
Shank position (posterior -> anterior)
h
120
80
40
50 ms
120
80
40
120
80
40
0
1
Power [a.u.]
i
Distal CA1/SUB stimulation
1
573
Unit #
-0.1
0
0.1
0.2
0.3
Time from stimulation onset [s]
5
4
3
2
1
0
-1
-2
-3
z-scored firing rate
j
Proximal CA1 stimulation
1
593
Unit #
-0.1
0
0.1
0.2
0.3
Time from stimulation onset [s]

**Supplementary Figure 5. Functional topology of hippocampal – gRSC connections in CaMKII-Cre::Ai32 animals.** a) Example histological verification of RSC targeting (left) as well as CA1 and subiculum targeting (right) using two 8-shank 256-site silicon probes coated with DiI (red). Dashed lines, L2/3 RSC. b) Exemplary voltage trace (top) and wavelet spectrogram averaged over 100 events (bottom) of iHFOs induced and recorded in the pyramidal cell layer of CA1. In all panels, the time of light stimulation is depicted by a blue bar. c) Z-score normalized raster plot showing the responses of CA1 units to CA1 light stimulation. d) Average LFP responses to >100 optogenetic stimulations from a proximal (blue), intermediate (green) and distal (red) shank. Response magnitude was comparable for intermediate and distal stimulation and slightly higher for proximal stimulation (mean ± s.e.m). e) Maximal RSC LFP response to light stimulation plotted against different stimulation intensities of proximal (blue) and distal (red) sites in dorsal CA1. Distal stimulation sites often included or bordered subiculum. Asterisks denote $p<0.001$. f) Maximal hippocampal LFP response (mean ± s.e.m) to light stimulation as a function of distance from stimulated shank from one example session. g) Normalized RSC LFP responses to >300 CA1 stimulations as a function of shank position along AP axis (mean ± s.e.m). h) Normalized wavelet spectrograms showing the spectral responses from a superficial RSC channel to a distal (top), intermediate (middle) and proximal (bottom) stimulation of CA1 and subiculum averaged over >300 events. i) Z-score normalized raster plot of RSC unit firing in response to distal CA1/ subicular stimulation sorted based on the location of maximal waveform amplitude (n = 3 sessions from 3 animals, targeting of superficial layers was histologically confirmed). j) Same, but for proximal CA1 stimulation.

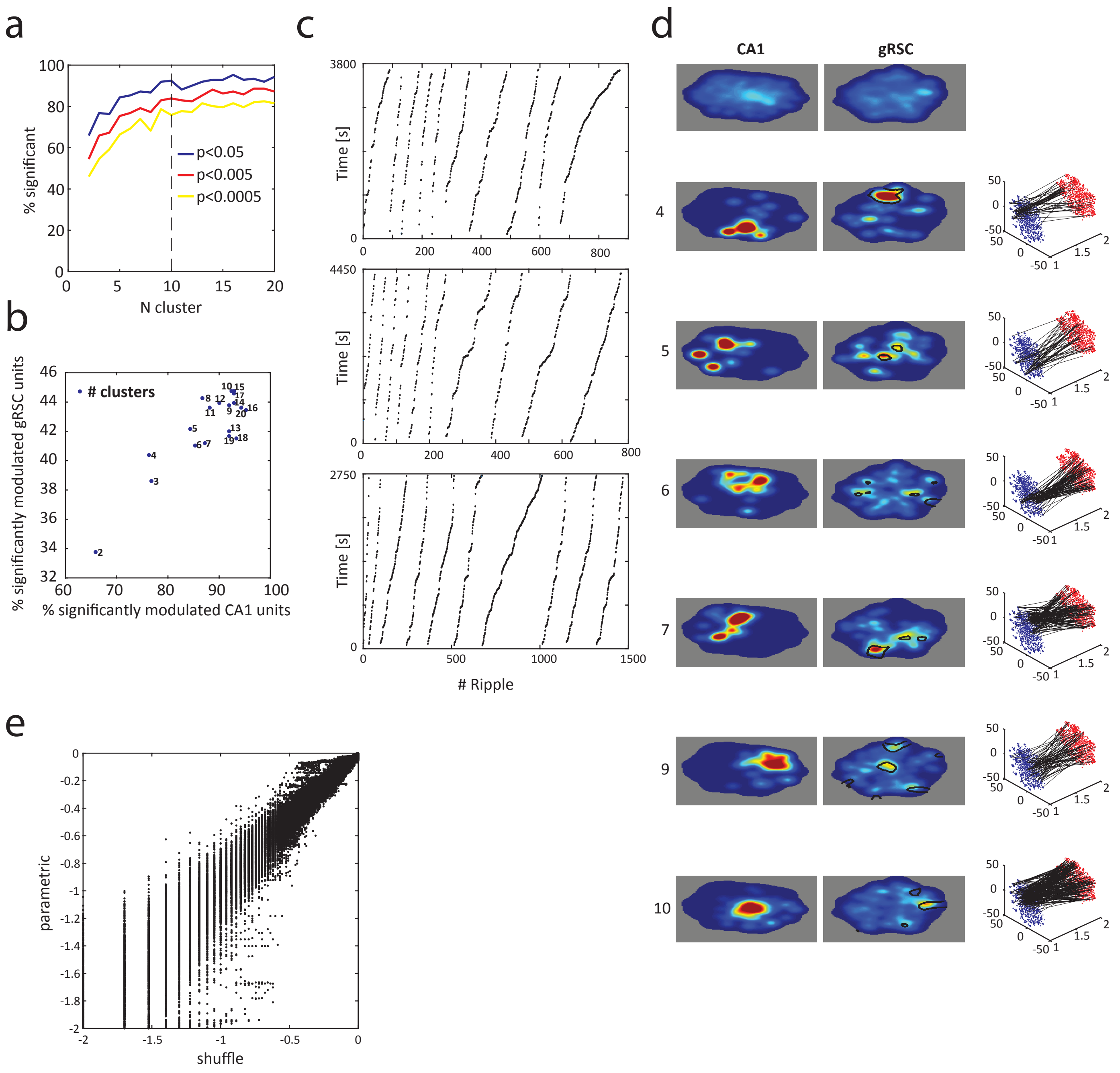


**Supplementary Figure 6. SPW-R clustering.** a) The percentage of CA1 neurons whose firing was significantly modulated by cluster ID (regressing out mean population firing rate), as a function of the number of clusters in the k-means analysis. Dashed line is set at k = 10, the number of ripple types used throughout. b) The percentage of neurons in CA1 and gRSC significantly modulated by ripple type for different number of clusters ($p < 0.05$). c) The distribution of SPW-Rs in time for each subject (n = 3), sorted by SPW-R cluster, showing that a SPW-R of a given type can occur at any point in time during the recording session. d) The t-SNE representation of each ripple type for one subject. Top panel shows occupancy plot for all ripples. See Fig. 5e. e) The correlation between the p-values obtained using a parametric ANCOVA test and after shuffling the labels of the ripple clusters and comparing the observed F-ratio from the ANCOVA test to the ones derived from the shuffle distribution.

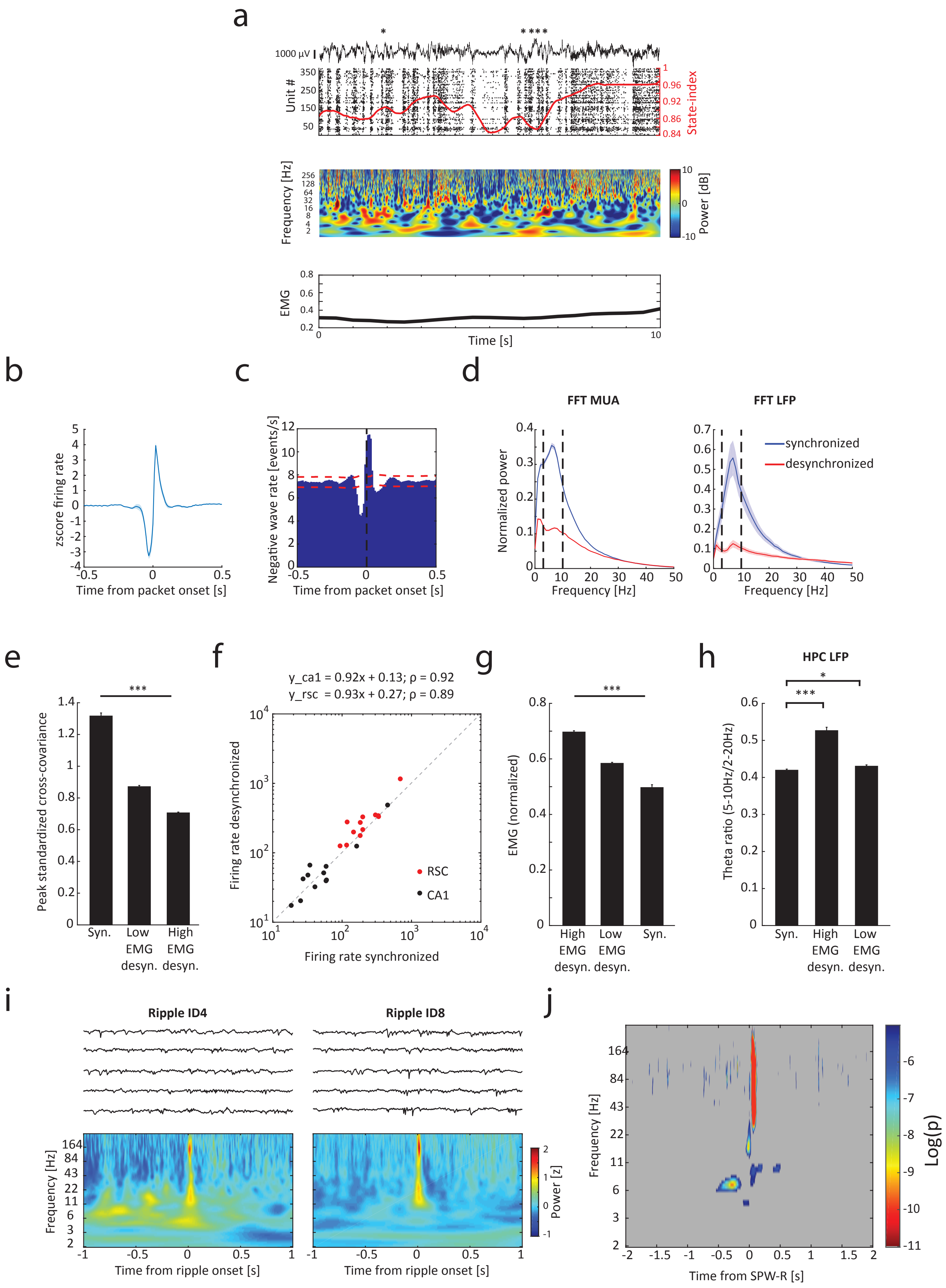
a
1000 µV
Unit #
State-index
Frequency [Hz]
Power [dB]
EMG
Time [s]
b
zscore firing rate
Time from packet onset [s]
c
Negative wave rate [events/s]
Time from packet onset [s]
d
FFT MUA
FFT LFP
Normalized power
Frequency [Hz]
synchronized
desynchronized
e
Peak standardized cross-covariance
Syn.
Low EMG desyn.
High EMG desyn.
f
y_ca1 = 0.92x + 0.13; ρ = 0.92
y_rsc = 0.93x + 0.27; ρ = 0.89
Firing rate desynchronized
Firing rate synchronized
RSC
CA1
g
EMG (normalized)
High EMG desyn.
Low EMG desyn.
Syn.
h
HPC LFP
Theta ratio (5-10Hz/2-20Hz)
Syn.
High EMG desyn.
Low EMG desyn.
i
Ripple ID4
Ripple ID8
Frequency [Hz]
Power [z]
Time from ripple onset [s]
j
Frequency [Hz]
Log(p)
Time from SPW-R [s]

**Supplementary Figure 7. Hippocampal-gRSC communication is state-dependent.** a) Top panel: example of 10s gRSC LFP (top), spiking activity (bottom) and the corresponding state-index (red) showing state transition from a synchronized to a desynchronized state. Asterisks, SPW-Rs detected in CA1. Middle panel: Wavelet spectrogram of the LFP trace shown above. Bottom panel: corresponding EMG activity. b) z-score normalized RSC MUA rate relative to packet onsets (n = 12 sessions from 4 animals, 58,987 packets). c) The rate of negative waves detected in superficial layers of the RSC relative to packet onset (n = 9 sessions from 4 animals 156,210 negative waves and 22,365 packets). d) Normalized power spectrum of MUA activity (left) and LFP (right) from concatenated epochs classified using the state index as either synchronized or desynchronized (8 sessions from 4 animals). e) Peak standardized cross-covariance (mean ± s.e.m) between CA1 and gRSC pairs of units from epochs classified as either synchronized or desynchronized, without or with increased EMG activity (n = 12 sessions from 5 animals 19,064 pairs). All groups are significantly different from each other ($p<0.001$ Bonferroni-corrected rank sum tests). f) A scatter plot showing the mean MUA rates in gRSC (red) and CA1 (black) across synchronized and desynchronized states defined using the state-index. Least-square slope and correlation coefficient for CA1 and gRSC are indicated above (n = 12 sessions from 5 animals). Dashed diagonal represents perfect correlation. Global MUA rates across synchronized and desynchronized states are unchanged in both structures (p's > 0.05, signed rank sum tests). g) EMG activity is significantly reduced during synchronized regime as compared to desynchronized states associated with either high or low EMG activity (p's < 0.001, Bonferroni corrected rank sum tests, n = 869 synchronized epochs, 1,526 desynchronized epochs with low EMG and 203 desynchronized epochs with high EMG ; 4 sessions from 4 animals). h) Narrow band theta expressed as the ratio between power at 5-10 Hz and 2-20 Hz in CA1 LFP is significantly lower in synchronized epochs, as compared to desynchronized epochs associated with high EMG ($p < 0.001$). Theta ratio of desynchronized epochs associated with low EMG activity is only marginally significantly different from that of synchronized epochs ($p = 0.046$, Bonferroni corrected rank sum tests). i) Additional examples of LFP traces (top) and averaged spectrograms for two ripple clusters. j) The degree to which gRSC power of each frequency band at each lag around ripple onset distinguishes CA1 population firing rate. This effect was regressed out from the degree of SPW-R type differentiation. Non-significant areas are shown in grey.

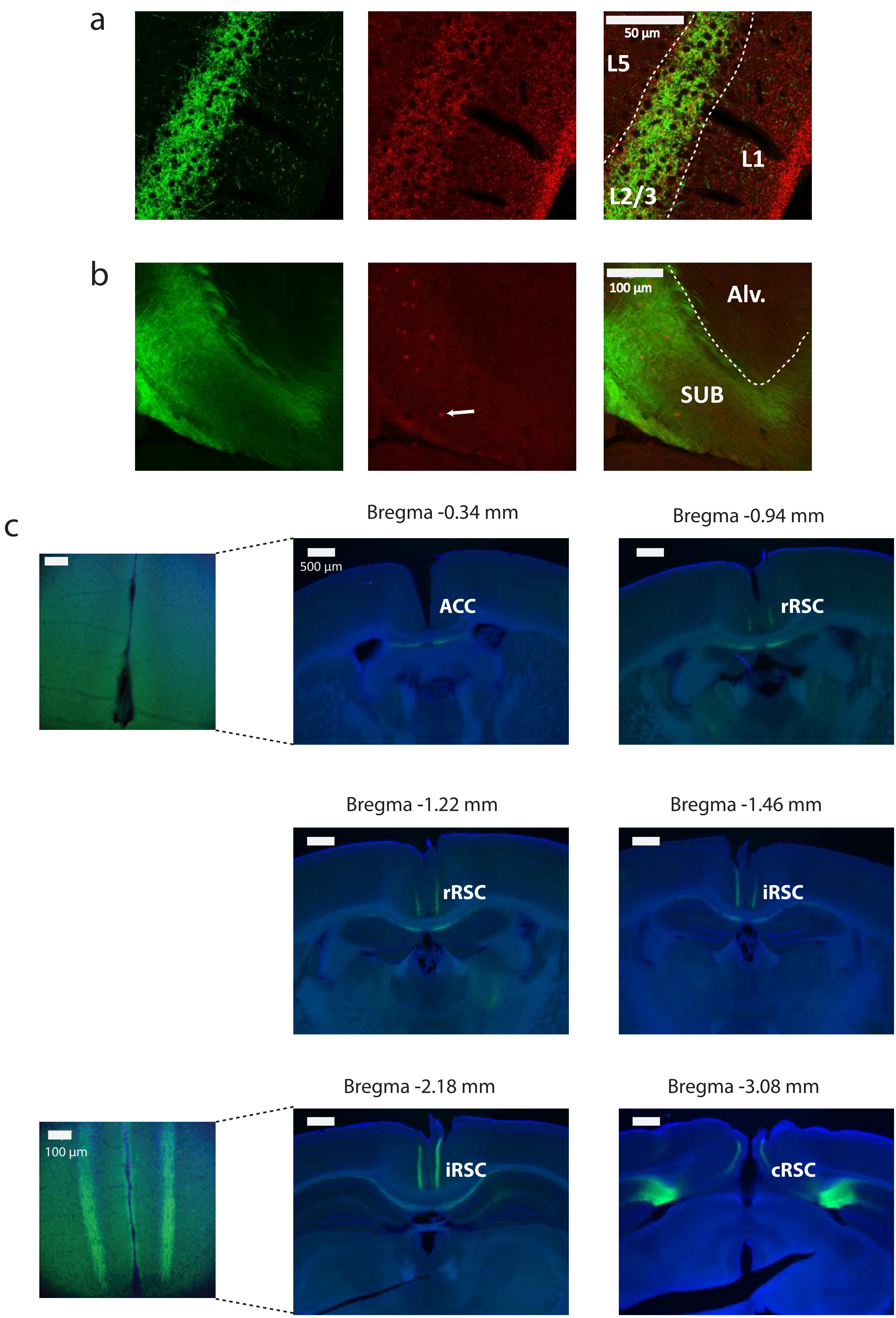
a
50 µm
L5
L1
L2/3
b
100 µm
Alv.
SUB
c
Bregma -0.34 mm
Bregma -0.94 mm
500 µm
ACC
rRSC
Bregma -1.22 mm
Bregma -1.46 mm
rRSC
iRSC
Bregma -2.18 mm
Bregma -3.08 mm
100 µm
iRSC
cRSC

**Supplementary Figure 8. Histology of subicular projections to gRSC in VGlu2-Cre animals.** a) VGlut2 immunostaining of brain slices from VGlut2-Cre animals injected with AAV9-SwitchON-mRubyN-LS-ChR2(H134R)-EYFP shows colocalization (right, merged) of subicular terminals (left, green) and VGlut2 antibodies (middle, red) in the superficial layers of the RSC (magnification: 60x). b) A Cre-switch leading to the expression of ChR2-eYFP (green) in Cre-positive cells and mRuby (red) in Cre-negative cells was used to assess the amount of virus spread from the area of injection and to exclude potential spread into the gRSC. White arrow depicts the most distal mRuby expressing cell detected in this section (magnification: 20x). Alv.: Alveus. c) Right: SUB – gRSC projections follow an anterior-posterior gradient showing strong axonal labeling (green) in caudal and intermediate sections (cRSC and iRSC, respectively) which gradually weakens in rostral sections of the RSC (rRSC) and terminates at the border with the ACC (magnification: 4x). Left images are 20x magnifications of ACC (top) and iRSC (bottom) sections. Blue: DAPI.

| | Rm (MΩ) | Vrest (mV) | Cm (pF) | AP threshold (mV) | Rheobase (pA) | AP height (mV) | AP FWHM (ms) | AHP (mV) | Sag (mV) | Tau (ms) |
|---|---|---|---|---|---|---|---|---|---|---|
| **L2/3 PYR** | 238.25 ± 9.91 | -79.26 ± 0.78 | 68.53 ± 6.22 | -35.36 ± 0.69 | 94.17 ± 4.66 | 66.54 ± 1.45 | 0.52 ± 0.01 | 20.52 ± 0.54 | -1.02 ± 0.09 | 15.41 ± 1.13 |
| **L5 PYR** | 140.10 ± 13.25 | -64.45 ± 1.05 | 272.10 ± 27.24 | -37.52 ± 1.07 | 115.00 ± 9.26 | 77.50 ± 2.64 | 0.60 ± 0.02 | 13.42 ± 0.87 | -11.85 ± 1.02 | 37.64 ± 3.28 |
| **FS INs** | 95.47 ± 13.98 | -73.40 ± 1.33 | 100.89 ± 18.97 | -37.27 ± 2.05 | 451.42 ± 105.27 | 65.28 ± 3.38 | 0.25 ± 0.01 | 24.62 ± 1.41 | -2.43 ± 0.64 | 8.13 ± 0.79 |

**Supplementary Table 1. Biophysical properties of RSC neurons.** Summary of biophysical properties (mean ± s.e.m.) of superficial pyramids (n = 44 cells), deep pyramids (n = 22 cells) and fast-spiking interneurons (FS INs; n = 7 cells, 5 from superficial and 2 from deep layers). Rm: membrane-resistance, Vrest: resting membrane potential Cm: capacitance, AP: action potential, AHP: after hyperpolarization, FWHM: full-width at half maximum.

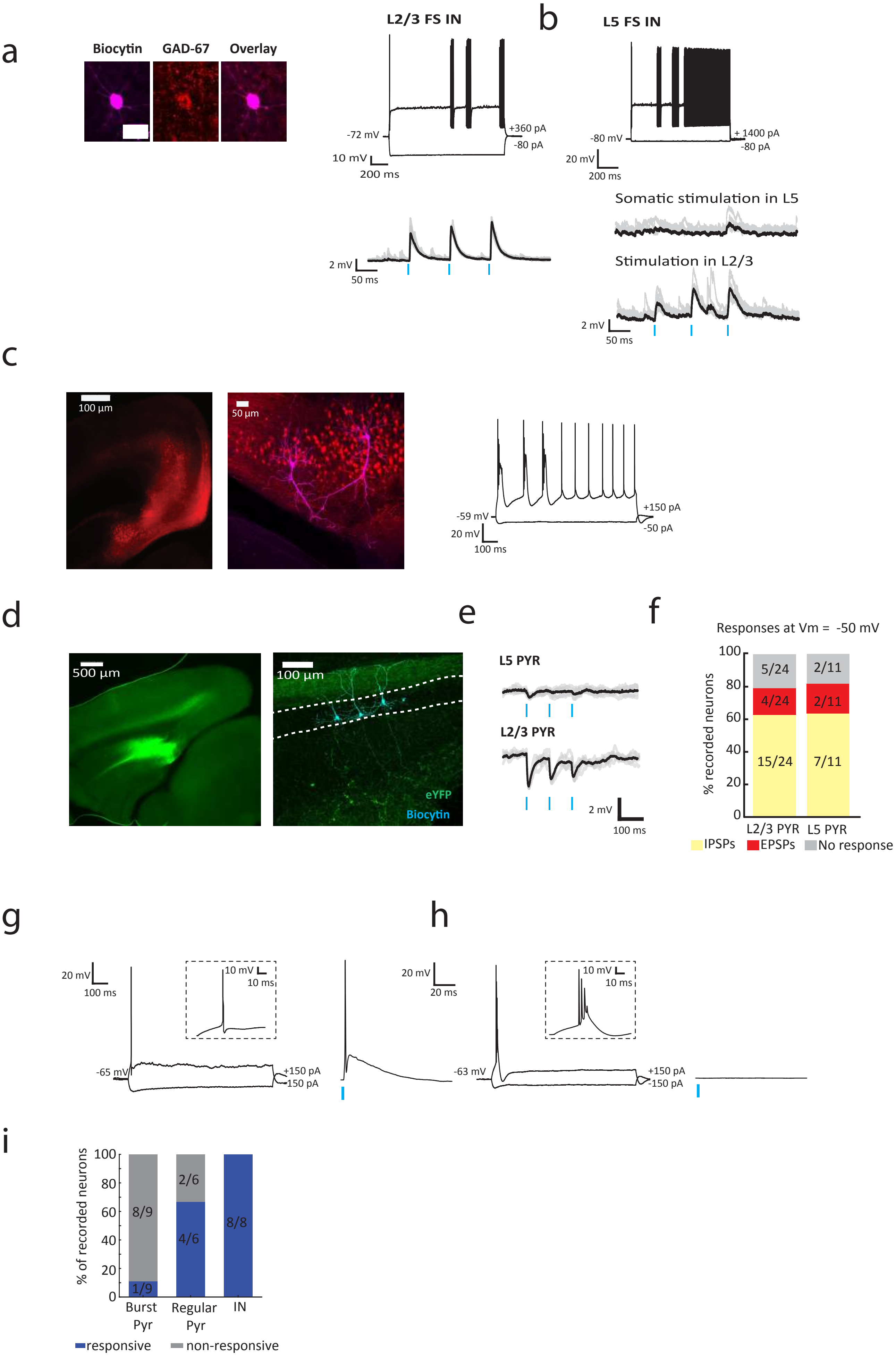

a
Biocytin
GAD-67
Overlay
L2/3 FS IN
-72 mV
+360 pA
-80 pA
10 mV
200 ms
2 mV
50 ms
b
L5 FS IN
-80 mV
+ 1400 pA
-80 pA
20 mV
200 ms
Somatic stimulation in L5
Stimulation in L2/3
2 mV
50 ms
c
100 µm
50 µm
-59 mV
+150 pA
-50 pA
20 mV
100 ms
d
500 µm
100 µm
eYFP
Biocytin
e
L5 PYR
L2/3 PYR
2 mV
100 ms
f
Responses at Vm = -50 mV
% recorded neurons
5/24
4/24
15/24
2/11
2/11
7/11
L2/3 PYR
L5 PYR
IPSPs
EPSPs
No response
g
20 mV
100 ms
10 mV
10 ms
-65 mV
+150 pA
-150 pA
h
20 mV
20 ms
10 mV
10 ms
-63 mV
+150 pA
-150 pA
i
% of recorded neurons
8/9
1/9
2/6
4/6
8/8
Burst Pyr
Regular Pyr
IN
responsive
non-responsive

**Supplementary Figure 9. Physiology of subicular projections to gRSC.** a) Example of a patched FS IN in superficial layers of the RSC, identified using GAD-67 immunostaining (left, magenta: biocytin, red: anti-GAD67 antibody), including firing characteristics (right, top) and example traces of light-evoked responses (right, bottom). b) FS INs in deep layers of gRSC exhibited light evoked responses which were stronger when stimulated in L2/3 (bottom trace) as compared to somatic stimulation in L5 (middle trace). Mean (black) and 10 exemplary traces (grey). Top, electrophysiological characterization. c) rAAV2-retro- mRuby injected in the RSC retrogradely labeled cells in distal subiculum (left, red). A subset of the labeled cells were patched (middle, magenta), all of which exhibited a bursty firing pattern (right, n = 8 cells from one animal). d) Left: a Cre-Off AAV-ChR2-YFP virus was injected into the dorsal subiculum to label VGlut2-negative cells. Right: subicular infection of VGlut2-Cre animals with Cre-off virus shows fiber labeling (green) predominantly in L1 and deeper layers, but to a lesser extent in L2/3. Three patched biocytin-filled pyramids are labeled in cyan. e) Examples of light-evoked responses from a deep (top) and a superficial (bottom) cell held at -50mV, revealing inhibitory inputs. f) Summary of light evoked responses from superficial and deep layers in VGlut2-Cre animals (n = 3 mice) injected with a Cre-Off AAV-ChR2-YFP virus. g) Electrophysiological firing pattern (left) and light-evoked responses (right) of an example subicular pyramidal cell infected by the Cre-Off AAV-ChR2-YFP virus, showing a regular firing pattern. Inset, zoomed-in action potential. h) Same, but for a subicular bursting cell. Note the absence of light-evoked responses. Same scale bars as in g. i) Summary of light-evoked responses in SUB cells (n = 23 cells from 3 animals).

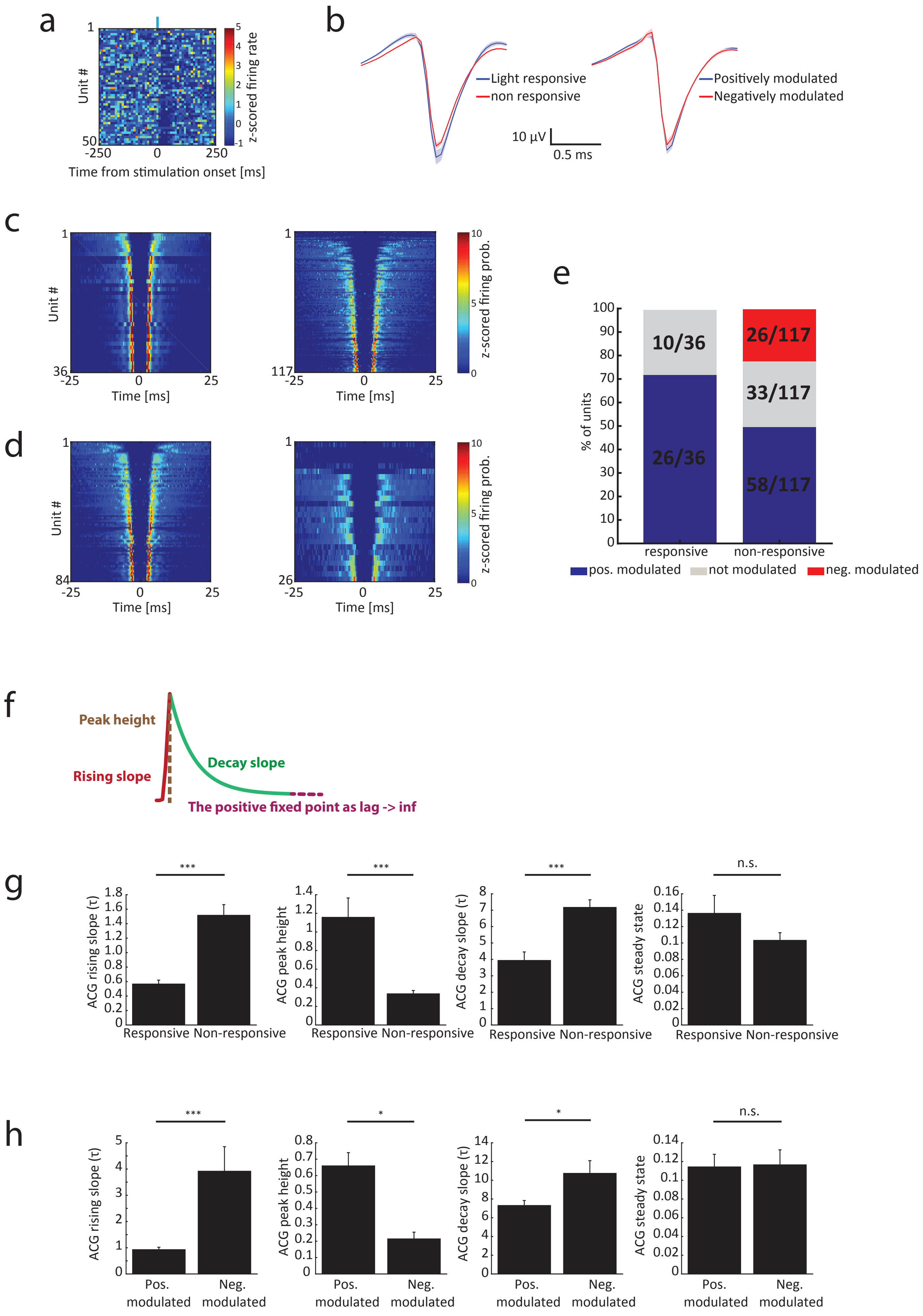
a
Unit #
Time from stimulation onset [ms]
z-scored firing rate
b
Light responsive
non responsive
Positively modulated
Negatively modulated
10 µV
0.5 ms
c
Unit #
Time [ms]
z-scored firing prob.
d
e
% of units
10/36
26/36
26/117
33/117
58/117
responsive
non-responsive
pos. modulated
not modulated
neg. modulated
f
Peak height
Rising slope
Decay slope
The positive fixed point as lag -> inf
g
ACG rising slope (τ)
ACG peak height
ACG decay slope (τ)
ACG steady state
Responsive
Non-responsive
n.s.
h
Pos. modulated
Neg. modulated

**Supplementary Figure 10. In vivo targeting of bursting subicular cells.** a) Optogenetic stimulation in VGlut2-Cre animals infected in the dorsal subiculum did not evoke neural firing in CA1 area (n = one session from one animal). b) Left: Average spike waveforms of light responsive (blue) and non-responsive (red) putative pyramidal cells. Right: Same but for SPW-R positively and negatively modulated putative pyramids (mean ± s.e.m). c) Color-coded z-scored ACGs of all light responsive (left) and non-responsive (right) gRSC units. d) Color-coded z-scored ACGs of all SPW-R positively (left) and negatively (right) modulated units. e) Summary of SPW-R modulation as a function of light responsiveness (n = 6 sessions from 3 animals; 36 light responsive units and 120 non-responsive units). f) A double exponential fit model of the ACG was used to determine significant differences in burstiness based on rising slope, peak height and decay slope. The positive lag reflects baseline firing probability. g) Summary of fit model parameters for light responsive and non-responsive units. h) Summary of fit model parameters for SPW-R positively and negatively modulated units (*$p<0.05$ ***$p<0.001$; rank sum tests).

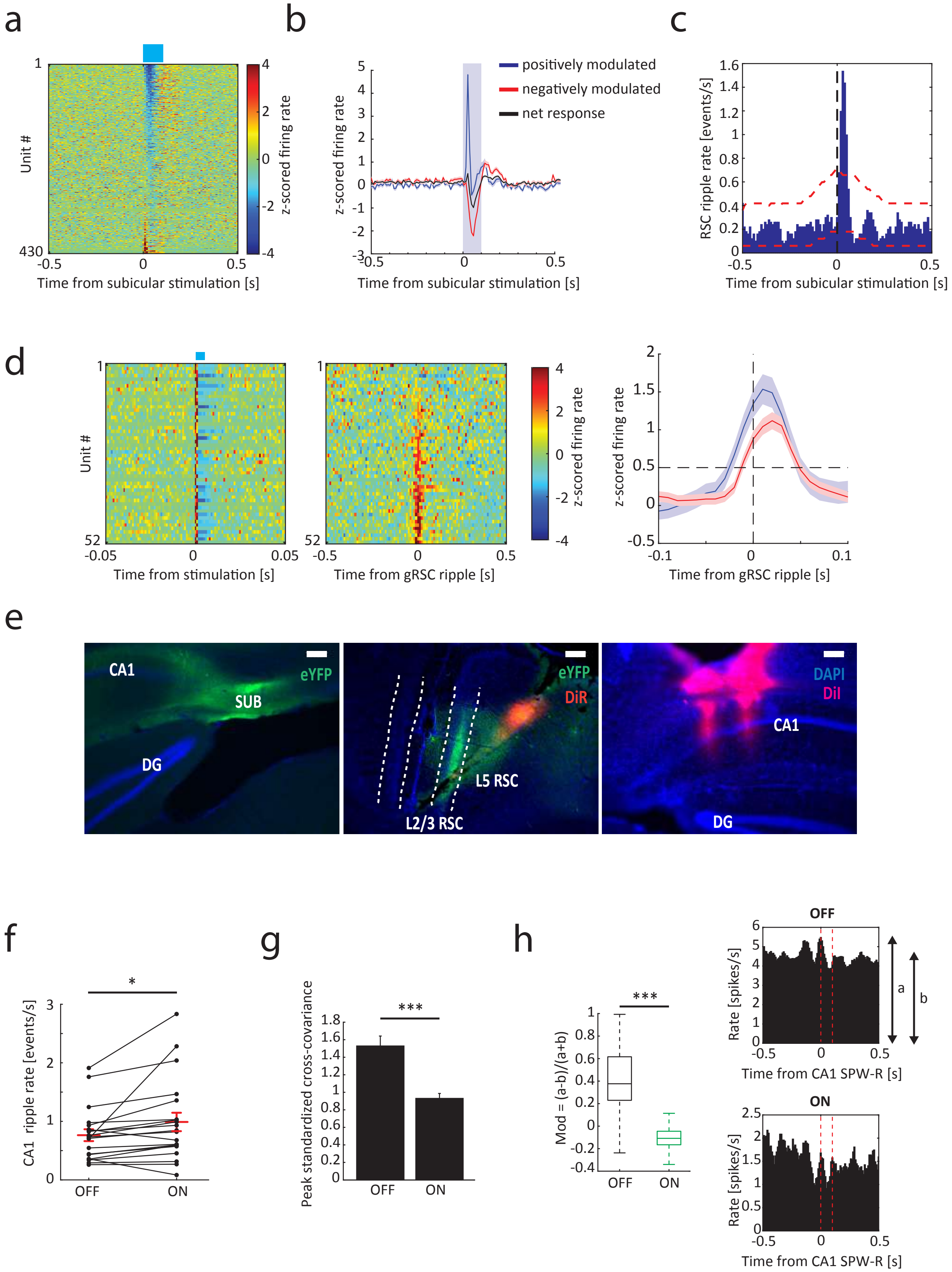

a
b
c
d
e
f
g
h
Unit #
z-scored firing rate
Time from subicular stimulation [s]
positively modulated
negatively modulated
net response
RSC ripple rate [events/s]
Time from stimulation [s]
Time from gRSC ripple [s]
CA1
SUB
DG
eYFP
DiR
L5 RSC
L2/3 RSC
DAPI
DiI
CA1 ripple rate [events/s]
OFF
ON
Peak standardized cross-covariance
Mod = (a-b)/(a+b)
Rate [spikes/s]
Time from CA1 SPW-R [s]

**Supplementary Figure 11. Subicular bursting neurons mediates SPW-R-related hippocampal output to the gRSC.** a) z-score normalized PETHs of gRSC units in response to 100 ms optogenetic square pulse stimulation of VGlut2+ cells in subiculum. b) Average of z-score normalized responses of units in gRSC which were positively (blue) and negatively (red) modulated by light stimulation, as well as net response (grey) (n = 15 sessions from 4 animals). c) Cross-correlation between subicular stimulation and identified gRSC ripple events (n = 7 sessions from 4 animals 1,672 sine wave stimulations and 3,718 gRSC ripples). d) Left: z-score normalized PETHs of putative pyramidal subicular cells which are significantly up-modulated by 5 ms blue light pulses (n = 7 sessions from 2 animals). Middle: z-score normalized PETHs of the same cells in response to gRSC ripples. Right: Average z-score normalized responses of the subicular cells shown in left and middle panels (blue) as well as of gRSC units (red, n = 175 units). Time to reach a z-score of 0.5 (dashed horizontal line) is significantly earlier for subicular cells (subicular cells: 26 ± 2 ms before gRSC ripple, gRSC cells: 15 ± 1 ms before gRSC ripple; p = 0.0009, rank-sum test). e) Arch-YFP expression in dorsal subiculum (left, green), as well as targeting of gRSC (middle) and hippocampal probes (right). f) CA1 SPW-R incidence is increased during stimulation of subicular VGlut2+ terminals in gRSC triggered on CA1 SPW-R detection (signed rank sum test, p = 0.003; n = 20 sessions from 4 animals;  28,220 ripples during light-off and 10,238 ripples during light-on). g) Peak standardized cross-covariance (mean ± s.e.m) between CA1 and gRSC unit pairs from spikes occurring ± 250 ms around CA1 SPW-Rs in either light-off or light-on periods (p = 0.0000002, rank sum test). h) Left: SPW-R modulation index (computed across all units which were significantly negatively modulated by Arch stimulation, n = 54), defined as the ratio between the peak of the SPW-R PETH between 0-100 ms after SPW-R onset (a) and a baseline period between -500 and -300 ms before SPW-R onset (b) is decreased for SPW-Rs occurring during light stimulation (p = 0.0001, z-statistics in shuffling analysis with 1000 shuffles). Right: PETH of an example unit for CA1 SPW-Rs occurring outside (top) or during (bottom) stimulation.
